# STUSC: selection tests under size change

**DOI:** 10.1101/2023.05.08.539872

**Authors:** Stephen Wooding

**Affiliations:** University of California Merced

## Abstract

**Summary:** Natural selection leaves signatures in genetic diversity. A challenge in detecting these signatures is that demographic history can shape genetic variation in ways similar to selection, making their effects difficult to disentangle. STUSC takes a simulation based approach to the problem. It performs five widely used tests for natural selection, Tajima’s D, Fu and Li’s F and F*, and Fu and Li’s D and D*, which ordinarily require the assumption of constant population size, under any size history. It also provides methods for generating their statistical distributions, allowing their responses to demography to be studied and past results to be evaluated.

**Availability and implementation:** STUSC is written in Java, and is freely available at the author’s website, woodinglab.org/stusc

## Introduction

Natural selection has strong effects on population genetic diversity. By favoring some alleles over others it alters their frequencies. Selection promoting an allele, for instance, can drive it rapidly to fixation. Conversely, selection against an allele can drive it to extinction. A requisite for these processes to occur is presence of functional variation; selection can only act on alleles with phenotypic effects. Thus, evidence of natural selection indicates the presence of functional polymorphism, its potential significance to organismal evolution, and its possible roles in inherited disease. A major objective in evolutionary biology is to find such evidence.

Theoretical approaches for detecting signatures of selection are well developed (Bamshad and Wooding, 2003; Vitti, et al., 2013). They capitalize the fact that selection alters multiple aspects of variation in genes under pressure. The number of polymorphisms, distribution of allele frequencies, levels of divergence among haplotypes, and other aspects of diversity are all affected. Tests for selection take advantage of this by comparing observed patterns of variation with expectations under selective neutrality, the null hypothesis. However, most tests are limited by an inability to take population size changes, which are common in natural settings and can mimic the effects of selection, into account. Population growth, for example, produces diversity patterns similar to those arising from positive selection, and population decline mimics the effects of diversifying selection. Thus, the effects of selection and demography are confounded, and difficult to separate.

The software described here, STUSC (Selection Tests Under Size Change) performs five tests widely used to detect signatures of selection in DNA sequence data: Tajima’s D, Fu and Li’s D and F, and Fu and Li’s D* and F* (Fu and Li, 1993; Tajima, 1989). These are both easily calculated and highly sensitive to the effects of selection, making them excellent tools for dissecting evolutionary processes. However, they rely on the assumption of constant population size, which limits their informativeness in many species of interest, including humans, whose numbers have increased exponentially over the last 100,000 years. Modeling the D and F statistics in dynamic populations can be done analytically but it is difficult except in special cases (Wooding and Rogers, 2002). However, simulating them is feasible. STUSC takes this approach, performing tests under any population size history specified by the user.

## Methods

STUSC simulates population genetic diversity under the coalescent model (Rosenberg and Nordborg, 2002). The coalescent focuses on gene genealogies. By defining the chances that pairs of individuals in any generation share a common ancestor in the previous one, the coalescent provides a probabilistic description of ancestor-descendant relationships. Population size changes are incorporated into the model by scaling levels of ancestor-descendant divergence up or down according to the population sizes at the time they occurred (Slatkin, 2001). This allows simulation of mutational processes, generating patterns of variation. Standard versions of the D and F tests determine expected patterns of diversity analytically for three measures sensitive to selective effects, the number of variable nucleotide positions (S), the number of nucleotide alleles observed just once in the sample (s), and the mean pairwise difference between sequences (mpd). To provide comparable tests, STUSC simulates the same measures.

STUSC takes three sets of parameters as input, which are detailed in the included documentation. First, it takes observed data needed to calculate statistics of interest to the user: the observed number of segregating sites (S, required to calculate all statistics), the mean pairwise difference between sequences (mpd, required for Tajima’s D and Fu and Li’s F and F*), and the number of singletons (s, required for all four of Fu and Li’s statistics). Next, it takes parameters describing an assumed population history, which is defined in an input file specifying changes in population size over time. Finally, STUSC takes parameters controlling the program’s execution and output, including the number of iterations to use for testing, formatting options, and verbosity.

STUSC specifies tests’ assumed population history under piecewise constant model. Under this model, history is represented as a series of one or more epochs, each defined by a size parameter, θ, and a time parameter, τ. θ (= 2N_e_μ) reflects the diversity expected if the epoch had a constant population size of N_e_, which is a function of the infinite sites mutation rate, μ. τ (=2μt) is a function of the time at which the epoch began in generations before present (t) and μ, which together mediate the amount of variation accumulated during that period. This structure allows any population size history to be analyzed (Rogers, 1995; Wooding and Rogers, 2002). For instance, a history of constant population size can be represented by one epoch, with a single θ and τ value, and populations experiencing incremental change, fluctuation, and other complex pasts can be represented by multiple epochs, each with a different θ and τ value. These are passed as input to STUSC in a comma separated value file, with each line containing the θ and τ representing an epoch, and epochs ordered from recent to ancient.

## Results

Example output from STUSC illustrates the impact of population history on the distribution of the D and F statistics (Figure 1). For instance, in a hypothetical population with an ancient θ = 10, the distribution of Tajima’s D is affected by both the magnitude of population growth and the time at which growth occurred, τ. Growth shifts the distribution in a negative direction proportional to the magnitude of growth, and τ shifts the distribution of D in a negative direction proportional to its closeness to θ (Figure 1A). Output under varying histories also illustrates how significance cutoff values for these tests are affected by population history (Figure 1B). For instance, the 5% cutoff in a sample of 1000 subjects with a θ of 10 and no population size change is D = 1.8, while the cutoff in the same population with 10-fold growth at τ = 10 has a 5% cutoff of D = -1.1, and the cutoff with 100-fold growth at τ = 10 has a 5% cutoff of D = -2.3.

**Figure 1.**
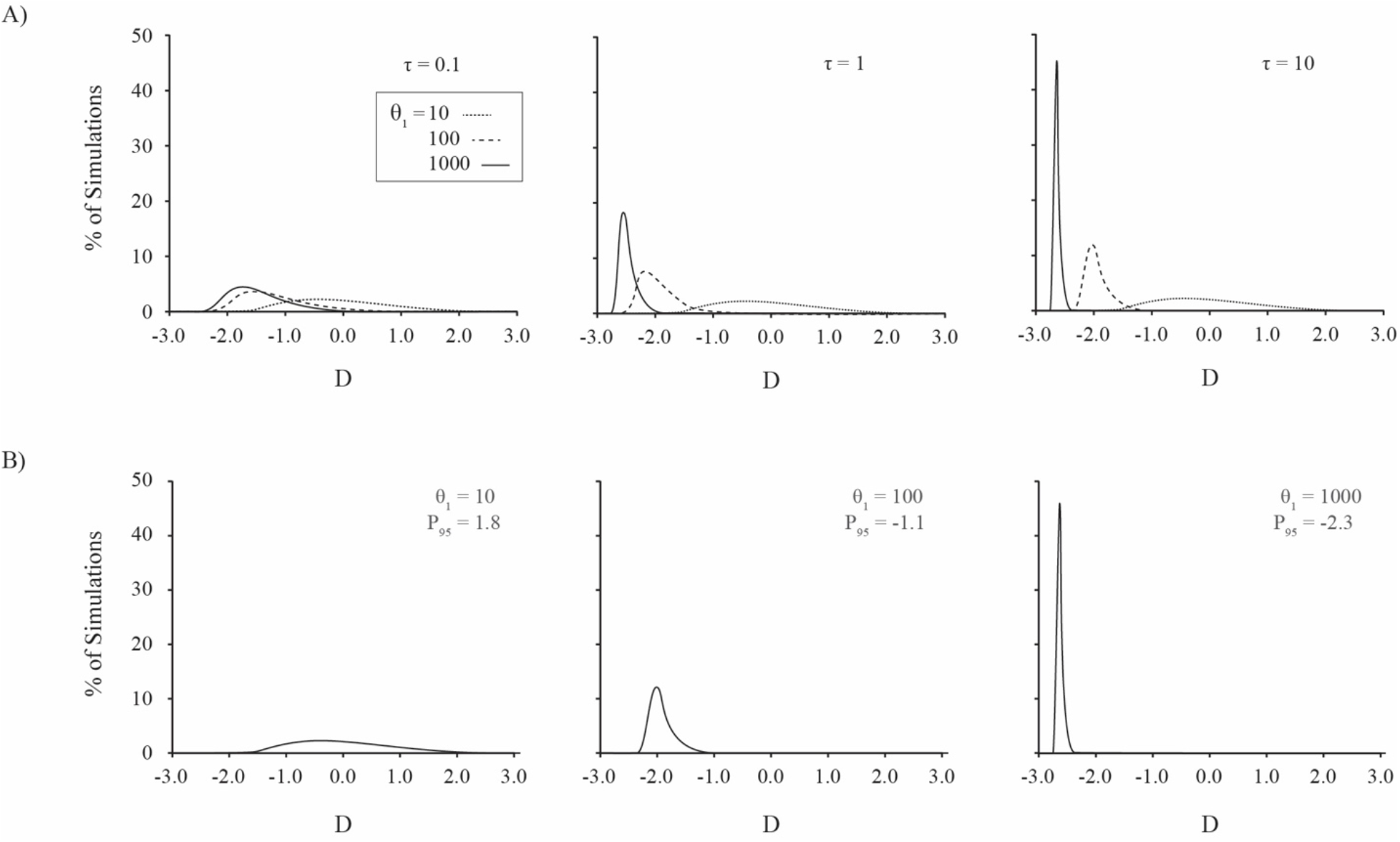
Effects of population growth on Tajima’s D statistic for two-epoch population histories. In each history θ_0_ denotes ancient θ, τ denotes time of population size change, and θ_1_ denotes modern θ. A) Distributions of D for a hypothetical sample of 1000 subjects with θ_0_ = 10, τ ranging from 0.1 to 10, and θ_1_ ranging from 10 to 1000. B) Effects of population growth on 5% significance cutoffs (P_95_) for D in a sample of 1000 subjects with θ_0_ = 10, τ = 10, and θ_1_ ranging from 10 to 1000.

## Conclusions

Population size change and natural selection are both common in natural populations, and both affect genetic diversity. In addition, their effects can be similar, making them difficult to separate. This hinders inferences about historic pressures and events based on genetic data; however, when population history is known, its effects can be subtracted, improving the detection of selective effects. STUSC provides a means for accomplishing this that is both flexible and simple to use.

